# Antagonistic Regulation of Circadian Output and Synaptic Development by the E3 Ubiquitin Ligase JETLAG and the DYSCHRONIC-SLOWPOKE Complex

**DOI:** 10.1101/769505

**Authors:** Angelique Lamaze, James E.C Jepson, Oghenerukevwe Akpoghiran, Kyunghee Koh

## Abstract

Circadian output genes act downstream of the clock to promote rhythmic changes in behavior and physiology, yet their molecular and cellular functions are not well understood. Here we characterize an interaction between regulators of circadian entrainment, output and synaptic development in *Drosophila* that influences clock-driven anticipatory increases in morning and evening activity. We previously showed the JETLAG (JET) E3 Ubiquitin ligase resets the clock upon light exposure, while the PDZ protein DYSCHRONIC (DYSC) regulates circadian locomotor output and synaptic development. Surprisingly, we find that JET and DYSC antagonistically regulate synaptic development at the larval neuromuscular junction, and reduced JET activity rescues arrhythmicity of *dysc* mutants. Consistent with our prior finding that DYSC regulates SLOWPOKE (SLO) potassium channel expression, *jet* mutations also rescue circadian and synaptic phenotypes in *slo* mutants. Collectively, our data suggest that JET, DYSC and SLO promote circadian output in part by regulating synaptic morphology.

**Highlights:**

- Loss of DYSC differentially impacts morning and evening oscillators
- Reduced JET activity rescues the *dysc* and *slo* arrhythmic phenotype
- Reduced JET activity causes synaptic defects at the larval NMJ
- JET opposes DYSC and SLO function at the NMJ synapse

## Introduction

Many aspects of adult behavior and physiology are subject to circadian regulation, including rhythmic alterations in locomotor activity that persist in the absence of changing environmental cues such as light or temperature. Forward genetic screens in the fruit fly, *Drosophila melanogaster*, have been of fundamental importance in elucidating the genetic underpinnings and molecular principles by which circadian clocks function and are entrained by the external environment (Allada and Chung, 2010; Axelrod et al., 2015; Franco et al., 2018; Tataroglu and Emery, 2015; Tategu et al., 2008). At the core of the *Drosophila* clock is a transcription-translation feedback loop, in which CLOCK (CLK) and CYCLE (CYC) activate transcription of their own inhibitors, PERIOD (PER) and TIMELESS (TIM). The molecular clock functions in approximately 150 neurons in the adult *Drosophila* brain (Nitabach and Taghert, 2008; Veleri et al., 2007). In standard 12h light-12h dark (LD) and constant 25°C conditions, *Drosophila* exhibit increased locomotor activity before light onset (morning anticipation) and offset (evening anticipation). These behavioral changes are driven by the circadian clock, and distinct groups of clock neurons in the lateral regions of the *Drosophila* brain, termed morning and evening oscillators, control morning and evening anticipation, respectively (Grima et al., 2004; Stoleru et al., 2004). However, whether the outputs of these distinct groups of clock neurons are regulated via the same molecular and circuit mechanisms is unclear.

Light is an important entraining factor for the *Drosophila* clock, and synchronizes circadian oscillations via degradation of TIM, thus re-setting the negative feedback component of the clock. This process is mediated by a molecular machinery comprised of the blue-light photo-receptor CRYPTOCHROME (CRY) and the F-box protein JETLAG (JET), a component of an E3 Ubiquitin ligase complex containing leucine rich repeats (Emery et al., 1998; Koh et al., 2006; Peschel et al., 2009; Stanewsky et al., 1998). Blue light activates CRY and induces the association of JET with both CRY and TIM, which facilitates proteasome-mediated degradation of TIM. To date, the sole known function for JET is in the light-induced TIM degradation pathway.

Several genes have been identified that, when mutated, result in arrhythmic locomotor patterns despite normal cycling of clock proteins (Dockendorff et al., 2002; Suh and Jackson, 2007; Williams et al., 2001), suggesting that these genes impact rhythmicity by regulating either the output of clock neurons or the function or development of downstream circuit components. We previously characterized a novel output gene termed *dyschronic* (*dysc*) (Jepson et al., 2012), a *Drosophila* homolog of *whirlin/DFNB31.* Human *DFNB31* encodes a PDZ-domain containing protein linked to Usher syndrome, a form of deaf-blindness (Mburu et al., 2003). *dysc* mutants exhibit arrhythmic locomotor behavior in constant dark (DD) conditions despite the persistence of wild-type oscillations in the molecular clock, and restoring *dysc* expression in clock neurons is not sufficient to rescue arrhythmicity (Jepson et al., 2012). This suggests DYSC functions in circuit components downstream of clock neurons for rhythmic behavior, but it may also function in clock neurons. DYSC binds to and regulates the expression of the calcium-activated voltage-gated potassium channel, SLOWPOKE (SLO), the *Drosophila* ortholog of the mammalian Slo1 BK potassium channel (Jepson et al., 2012). As with *dysc* mutants, loss-of-function mutations in *slo* result in arrhythmicity (Fernandez et al., 2007; Jepson et al., 2012). Furthermore, using the *Drosophila* larval neuromuscular junction (NMJ) as a model system, we demonstrated a novel role for DYSC in synapse development (Jepson et al., 2014). We found that DYSC is localized to synapses and closely associates with the active zone, the site of synaptic vesicle fusion. Loss of DYSC results in alterations in synaptic morphology, cytoskeletal organization, and active zone size (Jepson et al., 2014). Consistent with the regulation of SLO expression by DYSC, *slo* mutants exhibit NMJ phenotypes that overlap with those observed in *dysc* mutants. Together, these results demonstrate a role for DYSC and SLO in the regulation of circadian output and synaptic development.

Here we show that the morning and evening oscillators exhibit distinct functional requirements for DYSC under varying light conditions, and demonstrate an unexpected interaction between JET, DYSC and SLO in regulating both circadian rhythms and synaptic development. Mutations in *jet*, but not cry, significantly rescue the arrhythmic phenotype of both *dysc* and *slo* mutants in DD, suggesting a novel role for JET in the circadian output pathway. Intriguingly, synaptic defects in *dysc* and *slo* mutant synapses are also suppressed by *jet* mutations. We find that DYSC, SLO and JET act as regulators of synaptic bouton number and size, suggesting a common cellular locus for their activity. Collectively, these data raise the possibility that JET, DYSC and SLO may influence circadian output by regulating synaptic morphology.

## Results

### Loss of DYSC Differentially Impacts Morning and Evening Oscillators

Distinct clusters of clock neurons control rhythmic behavior under different environmental conditions (Grima et al., 2004; Stoleru et al., 2007; Stoleru et al., 2004; Zhang et al., 2010). We previously reported that *dysc* mutants are arrhythmic under DD conditions (Jepson et al., 2012). This indicates that DYSC is required for proper output of small lateral ventral neurons (s-LN_v_s) expressing the neuropeptide PDF, which controls rhythmicity in DD (Renn et al., 1999). PDF+ s-LNvs are referred to as the morning oscillator due to their role in anticipatory increase in locomotor activity before light onset (Grima et al., 2004). To examine the role of DYSC in additional clock neuron clusters, we assayed *dysc* mutants under different light conditions. As expected, *dysc* mutants did not exhibit morning anticipation in standard 12 h:12 h LD conditions (Figures 1A and S1A), confirming defective output of the morning oscillator. However, *dysc* mutants showed robust evening anticipation (Figures 1A and S1A), suggesting DYSC is not essential for the output of the CRY+, PDF-negative 5^th^ s-LN_v_ and dorsal lateral neurons (LN_d_s), referred to as the evening oscillator (Grima et al., 2004). We next examined locomotor behavior under prolonged photoperiod conditions (16 h: 8 h LD), in which evening anticipation is more readily visible as a peak of activity before light offset (Majercak et al., 1999; Schlichting et al., 2016; Yoshii et al., 2009). Consistent with the above data obtained in 12 h: 12 h LD, we observed clear evening peaks of activity in both wild type controls and *dysc* mutants under 16 h: 8 h LD (Figure 1B). These results suggest an essential role of DYSC for the output of the morning but not the evening oscillator in alternating light-dark conditions.

**Figure 1.**
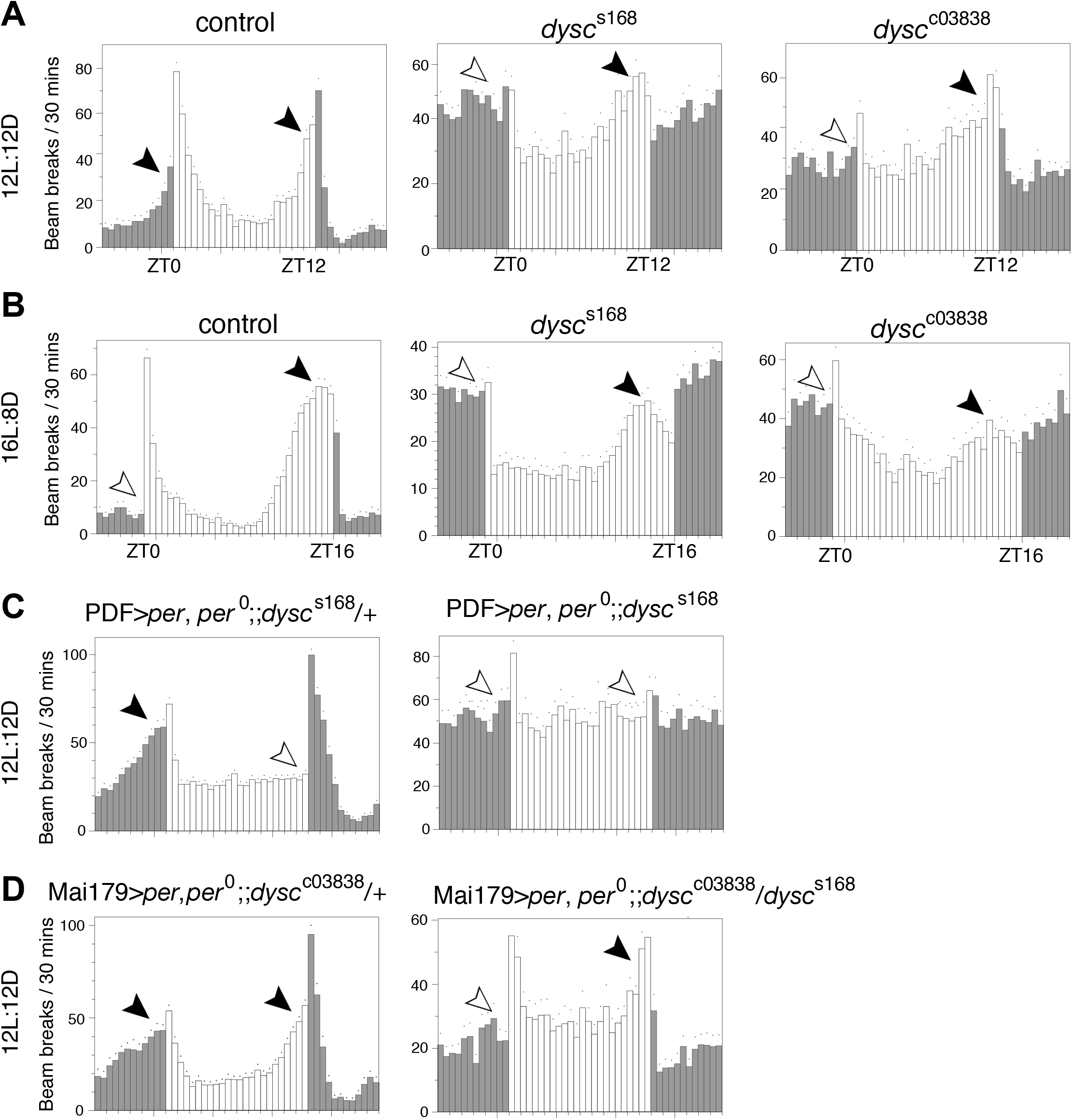
Loss of DYSC differentially impacts morning and evening oscillators. **(A)** Activity profiles of control and *dysc*^s168^ or *dysc*^c03838^ homozygous adult males in 12 h: 12 h light:dark conditions. Bars represent average beam breaks per 30 mins. Dots represent s.e.m. Gray bars represent dark periods, white bars light periods. In this and subsequent figures, filled and open arrowheads represent the presence and absence of anticipation, respectively. n = 38-54. Quantification of morning and evening anticipation is presented in Figure S1. **(B)** Activity profiles of males of indicated genotypes under extended photoperiod (16 h: 8 h light:dark) conditions. n = 28-53. **(C-D)** Activity profiles of *per*^0^ flies in which *per* is restored in the morning oscillator using PDF-Gal4 (C) or in both morning and evening oscillators using Mai179-Gal4 (D). The heterozygous *dysc* background (left) is used as a control for the *dysc* homozygous or transheterozygous (right) background. n = 24-66.

To further examine the differential role of DYSC in the output of the morning and evening oscillators, we analyzed the effects of restoring *per* expression in distinct groups of clock neurons of *per*^0^ mutants in *dysc* mutant versus control backgrounds. Previous studies have shown that expressing *per* only in the morning oscillator is sufficient to restore morning anticipation in LD, whereas expressing *per* only in the evening oscillator is sufficient to restore evening anticipation in LD (Grima et al., 2004; Stoleru et al., 2004). We compared the effects of restoring *per* expression in different group of clock cells in *per*^0^, *dysc*^s168^ double mutant flies vs *per*^0^, *dysc*^s168^/+ flies. We previously showed that *dysc* heterozygotes behave indistinguishably from wild-type control flies (Jepson et al., 2012), and as expected, restoring *per* in LN_v_s using PDF-Gal4 was sufficient to rescue morning anticipation in *dysc* heterozygotes. In contrast, it was not sufficient to rescue morning anticipation in *per*^0^, *dysc*^s168^ double mutants (Figures 1C and S1B), providing additional evidence that *dysc* is necessary for the output of the morning oscillator. Next, we used the Mai179-Gal4 driver to restore *per* expression in both the morning and evening oscillators (Grima et al., 2004). Whereas this was sufficient to rescue morning and evening anticipation in a *dysc* heterozygous background, only evening anticipation was rescued in a *dysc* homozygous background (Figures 1D and S1C). These results confirm that *dysc* is required for the output of the morning oscillator but not the evening oscillator in LD and suggest they may recruit non-overlapping downstream output pathways.

### Mutations in *jet*, but not cry, Rescue the *dysc* Arrhythmic Phenotype

Although *dysc* mutants exhibit clear evening anticipation, the pattern of anticipation is different from that of control flies. Whereas control flies rapidly increase their activity before the light offset over a few hours, *dysc* mutants gradually increase their activity over several hours (Figure 1A). To further examine the evening oscillator output in *dysc* mutants, we examined locomotor rhythmicity under different light conditions. In constant red light (RR), CRY is not activated and wild type flies maintain rhythmic behavior as do *cry* mutants in constant white light (LL). In LL, the evening oscillator drives rhythmic locomotor activity of *cry* mutants (Picot et al., 2007). Similarly, the evening oscillator drives rhythmicity in RR, as restoring *per* expression in both the morning and evening oscillator was sufficient to rescue arrhythmicity of *per*^0^ mutants, but restoring *per* in the morning oscillator alone was not (Figure S2). Since *dysc* mutants exhibited robust evening anticipation in LD conditions, we expected that they would be rhythmic in RR. However, we found that under RR conditions, *dysc* mutants were arrhythmic (Figures 2A and 2B), suggesting that the evening oscillator requires DYSC for its output in RR.

**Figure 2.**
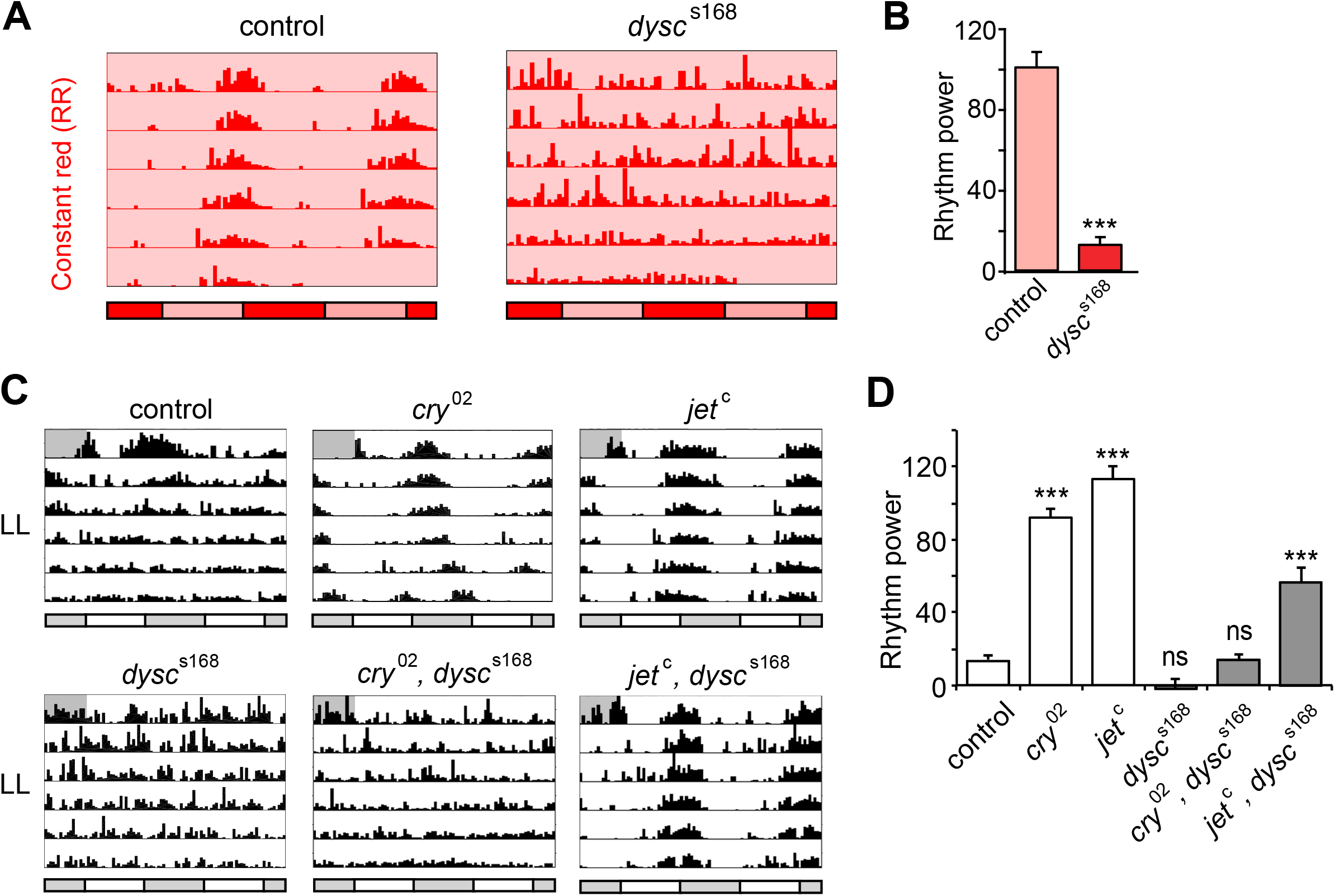
Reduced JET rescues arrhythmicity of *dysc* mutants in LL. **(A)** Representative actograms showing locomotor activity of control and *dysc*^s168^ homozygous males in RR, under which locomotor activity is driven by the evening oscillator (Figure S2). Red bars represent subjective night, pink bars subjective day. **(B)** Mean rhythm strength (power) in control and *dysc*^s168^ homozygotes under RR. n = 28-57. **(C)** Representative actograms of males of indicated genotypes in LL. **(D)** Mean rhythm strength (power) of males of indicated genotypes in LL. Whereas *dysc* single mutants are arrhythmic in LL, *jet, dysc* double mutants exhibit moderate rhythm strength. n = 24-66, except *dysc*^s168^, for which n = 14. Bars represent mean ± s.e.m. *** p < 0.0005, *t* test (B) and one-way Brown-Forsythe and Welch ANOVA for unequal variances followed by Dunnett T3 test relative to controls (D). Bars represent mean ± s.e.m.

We next examined the rhythmicity of *dysc* mutants in LL in a *cry* mutant background, which allows rhythmic behavior due to a functional evening oscillator (Picot et al., 2007; Stanewsky et al., 1998). Consistent with our above results in RR, double mutants for *dysc* and *cry* were arrhythmic in LL conditions (Figures 2C and 2D), further indicating that the evening oscillator is dependent on DYSC under constant light conditions.

Since CRY and JET act in the same pathway to regulate light-dependent TIM degradation and both *cry* and *jet* mutants are rhythmic in LL (Koh et al., 2006; Stanewsky et al., 1998), we expected *jet, dysc* double mutants to phenocopy cry, *dysc* double mutants in LL. Unexpectedly, however, we found that a hypomorphic mutation in *jet* (*jet*^c^) robustly restored rhythmicity to *dysc* mutants in LL (Figures 2C and 2D). To our further surprise, introduction of *jet*^c^ also rescued morning anticipation in *dysc* mutants under LD conditions, suggesting that reduced JET activity restores function to both the morning and evening oscillator output circuits in *dysc* mutants (Figures 3A and 3B). We observed morning anticipation using two independent alleles of *jet* (*jet*^c^ and *jet*^r^) and *dysc* (*dysc*^s168^ and *dysc*^c03838^) (Jepson et al., 2012; Koh et al., 2006). To further analyze morning oscillator output in *jet, dysc* double mutants, we next examined rhythmicity in DD. As in LL, we found a substantial (but not complete) rescue of locomotor rhythmicity in *jet, dysc* double mutants (Figures 3C and 3D). Together, these data demonstrate that reductions in JET activity partially rescue all known circadian phenotypes of the *dysc* mutant and reveal a novel function for JET in regulating circadian output that is independent of its canonical role in light entrainment.

**Figure 3.**
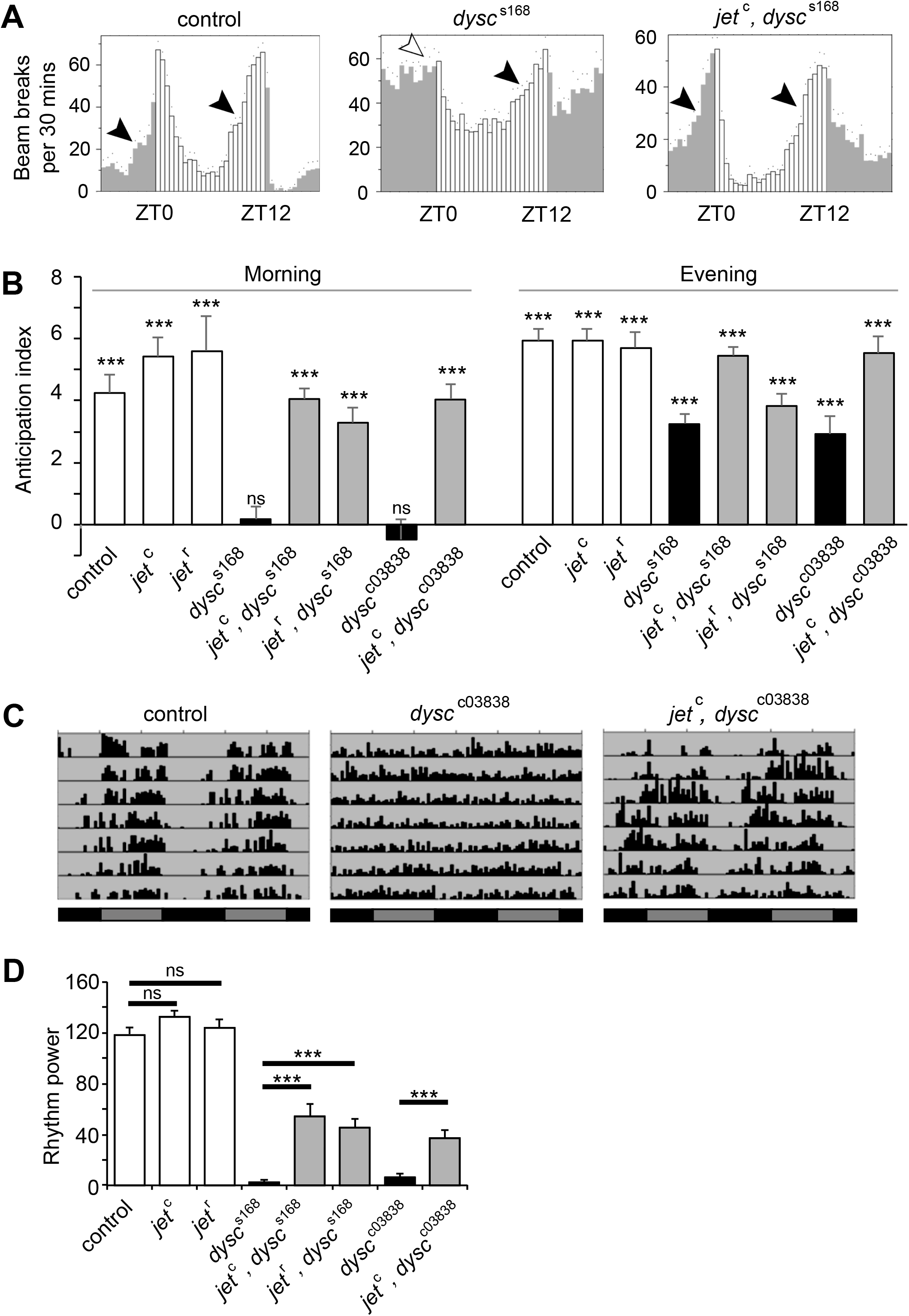
Reduced JET function restores morning anticipation in LD and rhythmicity in DD of *dysc* mutants. **(A)** Activity profiles of control males, as well as *dysc*^s168^ single and *jet*^c^, *dysc*^s168^ double mutants under 12 h: 12 h LD conditions. Morning anticipation, absent in *dysc* single mutants, is present in *jet, dysc* double mutants. **(B)** Morning and evening anticipation index for indicated genotypes. Anticipation index is defined as the slope of the best-fitting regression line for the activity counts over a period of 4 h (controls) or 8 h (all other genotypes) prior to a light-dark transition. n = 23-59. Bars represent anticipation index ± standard error, and statistics indicate whether anticipation index is significantly different from 0. **(C)** Representative actograms of control males, as well as *dysc*^c03838^ single and *jet*^c^, *dysc*^c03838^ double mutants under constant dark (DD). Black bars indicate subjective night, dark grey bars subjective day. **(D)** Mean power of rhythmicity of control adult males of indicated genotypes. n = 32-72. Bars represent mean ± s.e.m. *** p < 0.0005, ns: not significant, *t* test with Bonferroni correction (B); one-way Brown-Forsythe and Welch ANOVA for unequal variances followed by Dunnett T3 test for the indicated pairwise comparisons (D).

### Reduced JET Activity Rescues Circadian Phenotypes of *slo* Mutants

We previously showed that DYSC post-transcriptionally regulates the expression of the SLO BK potassium channel (Jepson et al., 2012). Mutations in *slo* also lead to arrhythmic locomotor behavior (Fernandez et al., 2007), suggesting that DYSC influences circadian output through its regulation of SLO. Since JET acts in the protein degradation pathway, a simple hypothesis is that reductions in JET activity protect SLO channels from proteasomal degradation following loss of DYSC, thus maintaining rhythmic behavior. This hypothesis yields a clear prediction that mutations in *jet* should be unable to rescue the arrhythmic phenotype of *slo* null mutants. We thus examined circadian function in *jet, slo* double mutants using flies harboring a P-element insertion in the second exon of the *slo* locus that acts as a null allele (*slo*^11481^) (Jepson et al., 2014). As with *dysc* mutants, *slo*^11481^ homozygotes were arrhythmic in DD (Figure 4A and 4B), and showed significant evening anticipation but no morning anticipation in LD (Figures 4C and S3). Intriguingly, in both DD and LD conditions, morning oscillator output was restored in *jet*^c^, *slo*^11481^ double mutants (Figures 4 and S3). Another simple hypothesis that DYSC itself is a substrate of JET is also unlikely because *jet* mutations rescue the circadian phenotypes of *dysc*^c03838^ null mutants. In addition, *dysc*^s168^ mutants express only a short-isoform of *dysc* that does not play a role in circadian rhythms and are essentially null for the isoforms that support rhythmic behavior (Jepson et al., 2012). Together, our data indicate that SLO and DYSC are not substrates of JET and demonstrate novel roles for JET that intersect with both DYSC and SLO to modulate circadian output.

**Figure 4.**
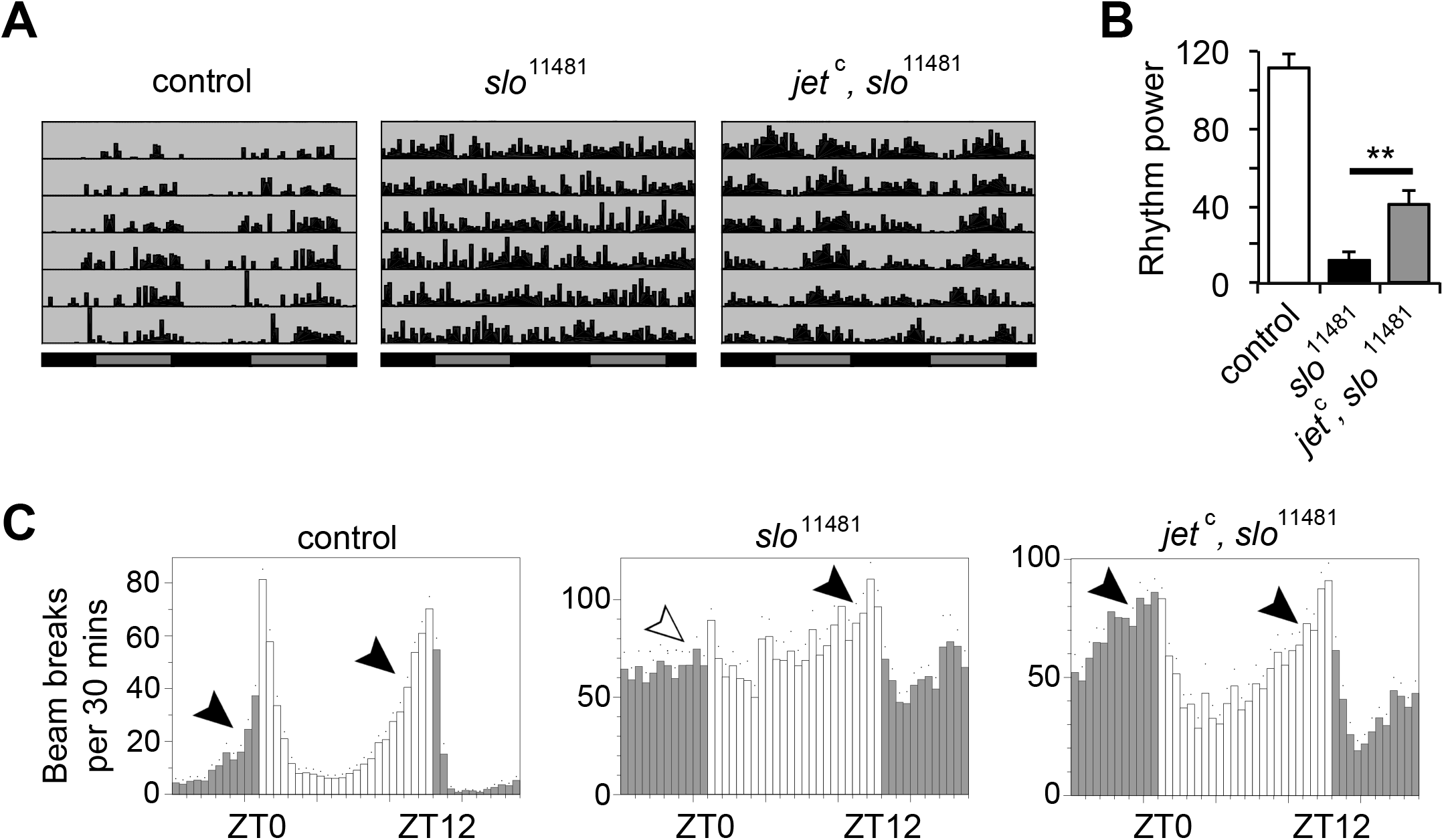
Reduced JET function restores morning anticipation in LD and rhythmicity in DD of *slo* mutants. **(A)** Representative actograms of control, *slo*^11481^ and *jet, slo*^11481^ males in DD. **(B)** Mean power of rhythmicity of control, *slo*^1148^ and *jet*^c^, *slo*^1148^ males in DD. n = 26-48. Bars represent mean ± s.e.m. ** p < 0.005, one-way Brown-Forsythe and Welch ANOVA for unequal variances followed by Dunnett T3 test for all pairwise comparisons, only one of which is shown for clarity. (C) Activity profiles of control males, as well as *slo*^11481^ single and *jet*^c^, *slo*^11481^ double mutants under 12 h: 12 h LD conditions. n = 35-43. Quantification of morning and evening anticipation is presented in Figure S3.

### Reduced JET Activity Rescues Synaptic Defects in *dysc* and *slo* Mutants

In the adult *Drosophila* brain, DYSC localizes to both neuronal tracts as well as the synaptic neuropil (Jepson et al., 2012), and at the larval neuromuscular junction (NMJ), DYSC exhibits a punctate pre-synaptic expression pattern in synaptic boutons and localizes closely to active zones, the sites of neurotransmitter release (Wagh et al., 2006). We previously demonstrated that DYSC plays an important role in modulating a range of synaptic parameters, with loss of DYSC resulting in a reduction in the number of synaptic boutons coupled with an increase in bouton size (Jepson et al., 2014). Thus, we were interested to determine whether reduced JET function could modify the synaptic effects of loss of DYSC.

To do so, we examined synaptic morphology at the 3^rd^ instar NMJ in wild type larvae compared to *jet* and *dysc* single mutant larvae and *jet, dysc* double mutants. Consistent with our previous data, loss of DYSC resulted in a significant reduction in synaptic bouton number and enlarged boutons (Figure 5). Interestingly, reduced JET activity resulted in a marked increase in bouton number, revealing a previously uncharacterized role for JET in synaptic development (Figures 5A and 5B). As expected from the opposite effects on bouton number, reduced JET compensated for the loss of DYSC, and *jet, dysc* double mutants had essentially wild-type number of boutons (Figures 5A and 5B). Intriguingly, despite the fact that reduced JET activity resulted in a slight but significant increase in bouton size, it nonetheless partially rescued the much greater increase in bouton size of *dysc* mutants (Figures 5A and 5C), demonstrating an interaction between the two genes for the regulation of bouton size. As expected from the similarities between *dysc* and *slo* bouton size phenotypes, reduction of JET activity also rescued the large bouton size phenotype of *slo* mutants (Figures 5A and 5C). As previously shown, unlike *dysc* mutants, *slo* mutants show a wild-type number of boutons. Interestingly, loss of SLO can rescue the increased bouton number phenotype of *jet* mutants (Figures 5A and 5B), further demonstrating opposing roles of *slo* and *jet* for synapse development. Thus, JET acts in opposition to DYSC and SLO in a common pathway regulating synaptic morphology.

**Figure 5.**
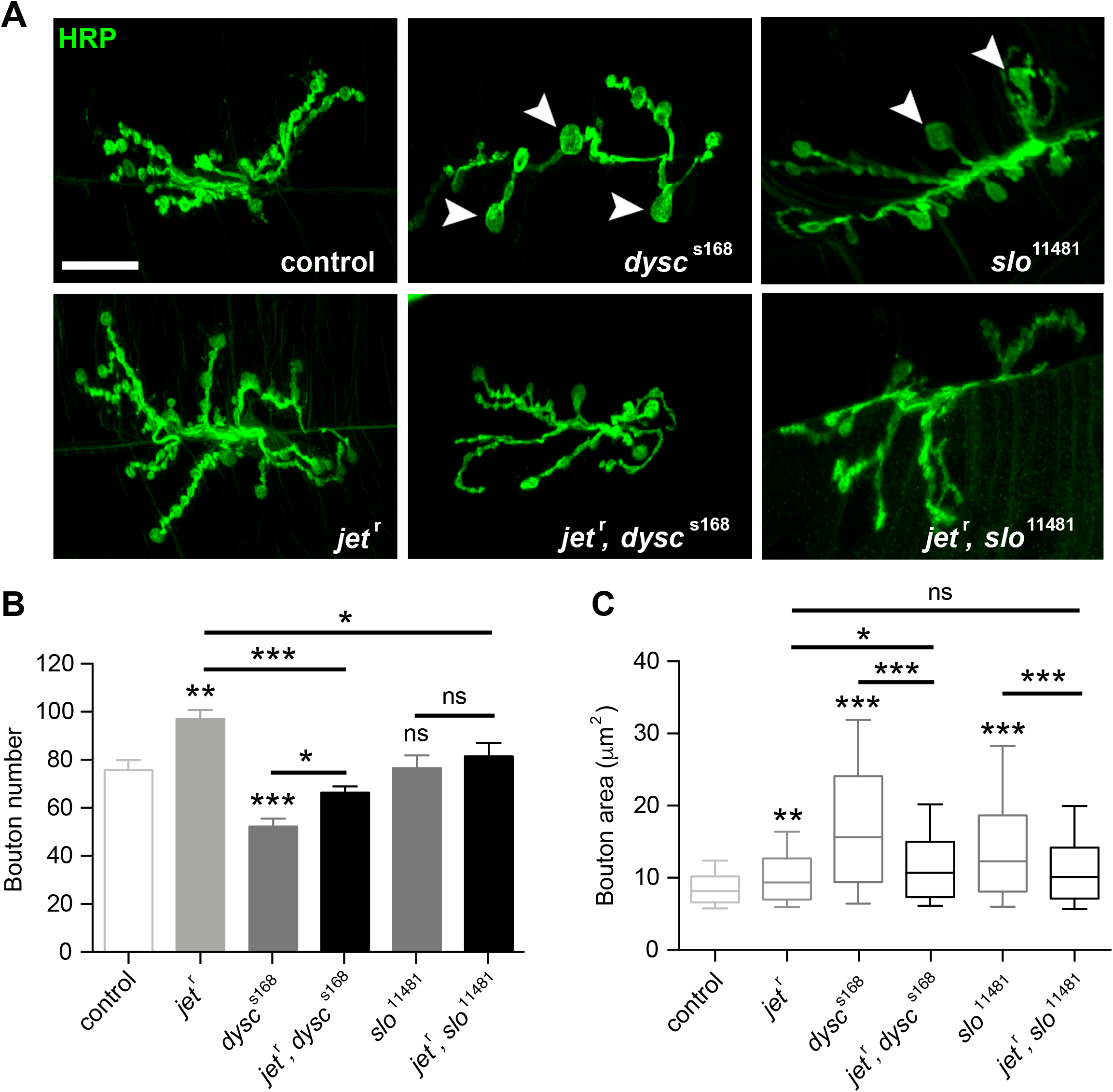
Mutations in *jet* rescue synaptic defects in *dysc* and *slo* homozygous larvae. **(A)** Representative confocal z-stacks showing HRP-labeled synapses from muscle 6/7, segment 3, of the NMJ of the 3^rd^ instar larvae. Note the enlarged boutons in *dysc*^s168^ and *slo*^11481^ mutant synapses (arrowheads), which are no longer present in *jet*^r^, *dysc*^s168^ and *jet*^r^, *slo*^11481^ double mutant synapses. Scale bar, 20 μm. **(B)** Average number of synaptic boutons in control, *jet*^r^, *dysc*^s168^, *slo*^11481^ single and double mutant synapses. Bars represent mean ± s.e.m. n = 16-30. **(C)** Box plot illustrating distribution of synaptic bouton areas in the indicated genotypes. Whiskers/lines represent 10^th^, 25^th^, median, 75^th^, and 90^th^ percentiles of the population distribution. n = 188-391. * p < 0.05, ** p < 0.005, *** p < 0.0005, ns: not significant, one-way Brown-Forsythe and Welch ANOVA for unequal variances followed by Sidak post-hoc test (B), and Kruskal-Wallis test followed Dunn post-hoc test (C). For both (B) and (C), 7 pairwise post-hoc tests were performed to compare controls against single mutants and double mutants against single mutants.

## Discussion

The present investigation began with the interesting observation that whereas *dysc* mutants are arrhythmic in DD and lack morning anticipation in LD conditions, they show robust evening anticipation. This led us to hypothesize that DYSC is required for the proper functioning of the morning oscillator but not of the evening oscillator. We found that although the evening oscillator in *dysc* mutants is functional in LD conditions, it is unable to support rhythmic behavior in RR or LL conditions. A plausible interpretation is that the loss of DYSC causes mild defects in the output of the evening oscillator, which have a relatively minor impact on behavior in oscillating LD conditions where the clock is reset on a daily basis, but have a stronger impact on rhythmic behavior over several days in constant conditions.

Examination of *dysc* mutants in *cry* or *jet* mutant background revealed a novel role of JET distinct from its known role for light entrainment of the clock. Although *jet* mutants do not exhibit any defects in rhythmicity in DD conditions, the fact that *jet* mutations can rescue *dysc* phenotypes suggests that JET plays a role in maintaining rhythmic behavior in constant conditions. Furthermore, *jet* mutants exhibit an increase in synaptic bouton number at the NMJ, providing direct evidence of a role for JET in synaptic development.

DYSC and SLO bind to each other and regulate each other’s expression, and *dysc* and *slo* mutants have similar circadian and synaptic phenotypes (Jepson et al., 2012; Jepson et al., 2014). Thus, it is not surprising that *jet* mutations can also rescue *slo* mutant phenotypes. Not only can *jet* mutations rescue *dysc* and *slo* circadian and enlarged bouton phenotypes, *dysc* and *slo* mutations can also rescue the increased bouton number phenotype of *jet* mutants. This highlights the antagonistic nature of the interaction between JET and the DYSC-SLO complex.

Our finding that *jet* mutations can rescue null mutations of *dysc* and *slo* indicate that DYSC and SLO are not substrates of JET. Thus, an important future goal would be to identify the substrates of JET relevant for circadian output and synaptic development. This in turn may help us to understand how JET, DYSC, and SLO interact and whether the same substrates mediate their roles in circadian output and synaptic development.

The fact that all three molecules function in both circadian rhythms and synaptic development suggest that the two processes are closely linked. Clock neurons as well as downstream circuit components in *dysc* and *slo* mutants may have synaptic defects analogous to those observed at the NMJ, which may mediate their circadian phenotypes. Similarly, compensatory synaptic alterations in clock and downstream neurons caused by *jet* mutations may underlie their ability to restore rhythmicity in *dysc* and *slo* mutants. It would be fruitful to investigate whether the antagonistic functions of JET vs DYSC and SLO during larval synaptic development have counterparts in the adult brain, especially in clock neurons.

In summary, our research reveals novel roles of JET in opposition to DYSC and SLO for circadian behavior and synaptic development. The functions of the closest mammalian homolog of JET, F-box and LRR Protein 15 (FBXL15), in circadian rhythms and synaptic development are unknown. It will therefore be intriguing to examine whether FBXL15 and the mammalian homologs of DYSC and SLO, Whirlin and Slo1 respectively, similarly function in an antagonistic manner to regulate circadian rhythms and synaptic development.

## Methods

### Fly strains

Flies were raised on standard food containing cornmeal, yeast, and molasses. Isolation of *jet*^c^, *jet*^r^, and *dysc*^s168^ was previously described; *cry*^02^ was obtained from Charlotte Helfrich-Förster (Dolezelova et al., 2007; Gegear et al., 2008); *per*^0^ and UAS-*perl6* from François Rouyer (Blanchardon et al., 2001); *dysc*^c03838^ from the Exelixis collection at the Harvard Medical School; and *slo*^11481^ from the Bloomington Stock Center (BL29918). PDF-Gal4 and Mai179-Gal4 (Grima et al., 2004; Siegmund and Korge, 2001) were obtained from Amita Seghal and François Rouyer, respectively.

### Locomotor assays

To assay locomotor behavior, 2- to 5-day old male flies were entrained to a 12 h: 12 h LD cycle for at least 3 days and placed individually in glass tubes containing 5% sucrose and 2% agar. Locomotor activity was monitored using the *Drosophila* Activity Monitoring System (Trikinetics) at 25°C under indicated light conditions. For quantification of circadian behavior in constant conditions (DD, RR or LL), activity counts (beam breaks) were collected in 30-min bins over a 6-day period, and power of rhythmicity was determined using Fly Activity Analysis Suite for Mac OSX (FaasX, M. Boudinot). The power of rhythmicity is defined as the difference between the *χ*^2^ value and the significance value at *p*=0.05. Average rhythm power was determined for all flies including arrhythmic ones. Individual actograms were generated using ClockLab (Actimetrics), and 24 h activity plots showing the mean beam breaks per 30-min bin over a 1-3 day period were generated using FaasX, which was also used to calculate anticipation index. Anticipation index is defined as the slope of the best-fitting linear regression line over a period of several hours prior to the light-dark transition. To account for the differences in the duration of anticipatory behavior, a period of 4 h was used for control flies and *jet* mutants, and 8 h for all other genotypes. The slope was normalized so that the maximum activity count for a given period was set to 50 (for 4 h periods) or 100 (for 8 h periods). This normalization procedure yields comparable anticipation indices for activity increases of similar magnitudes independent of the duration of anticipation.

### Measurement of synaptic bouton number and size at the larval NMJ

The morphology of synapses from muscle 6/7, segment 3, from wandering 3^rd^ instar larvae was assessed using an Olympus Fluoview confocal microscope. Larvae from both sexes were pooled. Larvae were dissected in a low Ca^2+^ HL3.1 solution of composition (mM): 70 NaCl, 5 KCl, 0.2 CaCl_2_, 20 MgCl_2_, 10 NaHCO_3_, 5 Trehalose, 115 Sucrose, 5 HEPES, pH 7.2. Dissected larval fillets were fixed in 4% PFA (in PBS) for 10-20 mins at room temperature. For immuno-staining, antibodies were diluted in PBS with 0.15% Triton-X and 5% goat serum. Fluorescently conjugated (Alexa-488) anti-HRP (Jackson ImmunoResearch, 1:200) was used to label neuronal membranes. The number of boutons was quantified for each synapse using an antibody against cysteine string protein (CSP; 6D6-c; 1:1000) (Developmental Studies Hybridoma Bank), which specifically labels the presynaptic bouton (Zinsmaier et al., 1990). Goat anti-mouse Alexa-fluor 555 (Molecular Probes, 1:1000) was used to label the anti-CSP primary antibody. Quantification of synaptic morphological parameters was performed and analyzed blind to genotype. Type 1b and Type 1s boutons were counted simultaneously. Synaptic bouton size was measured using ImageJ, with a minimum cut-off size of 5 μm^2^ (Jepson et al., 2014).

### Statistical Analysis

GraphPad Prism was used for statistical tests. Either *t* tests for pairs of groups or one-way ANOVAs for multiple groups were performed. If the groups had unequal variances, *t* tests for unequal variances or the Brown-Forsythe version of ANOVA was used. ANOVAs were followed by Dunnett, Tukey, or Sidak post-hoc tests depending on the number and type of pairwise comparisons performed. For comparisons of non-normally distributed data, Kruskal-Wallis tests were performed, followed by Dunn’s post-hoc tests.

## Acknowledgements

We thank Drs. Charlotte Helfrich-Förster, Francois Rouyer, and Amita Seghal, and the Bloomington Stock Center for fly stocks; M. Boudinot and Dr. Francois Rouyer for the FaasX software; and Huihui Pan for technical assistance. This work was supported by grants from the National Institutes of Health (R01GM088221 and R01NS086887 to K.K.).

## Author Contributions

A.L., J.E.C.J., and K.K. conceived the study. A.L., J.E.C.J., performed experiments and analyzed data with the assistance of O.A. A.L., J.E.C.J., and K.K. wrote the paper and O.A. commented on it. K.K. provided funding and supervision.

