## Supplemental Figures for "Antagonistic Regulation of Circadian Output and Synaptic Development by the E3 Ubiquitin Ligase JETLAG and the DYSCHRONIC-SLOWPOKE Complex"

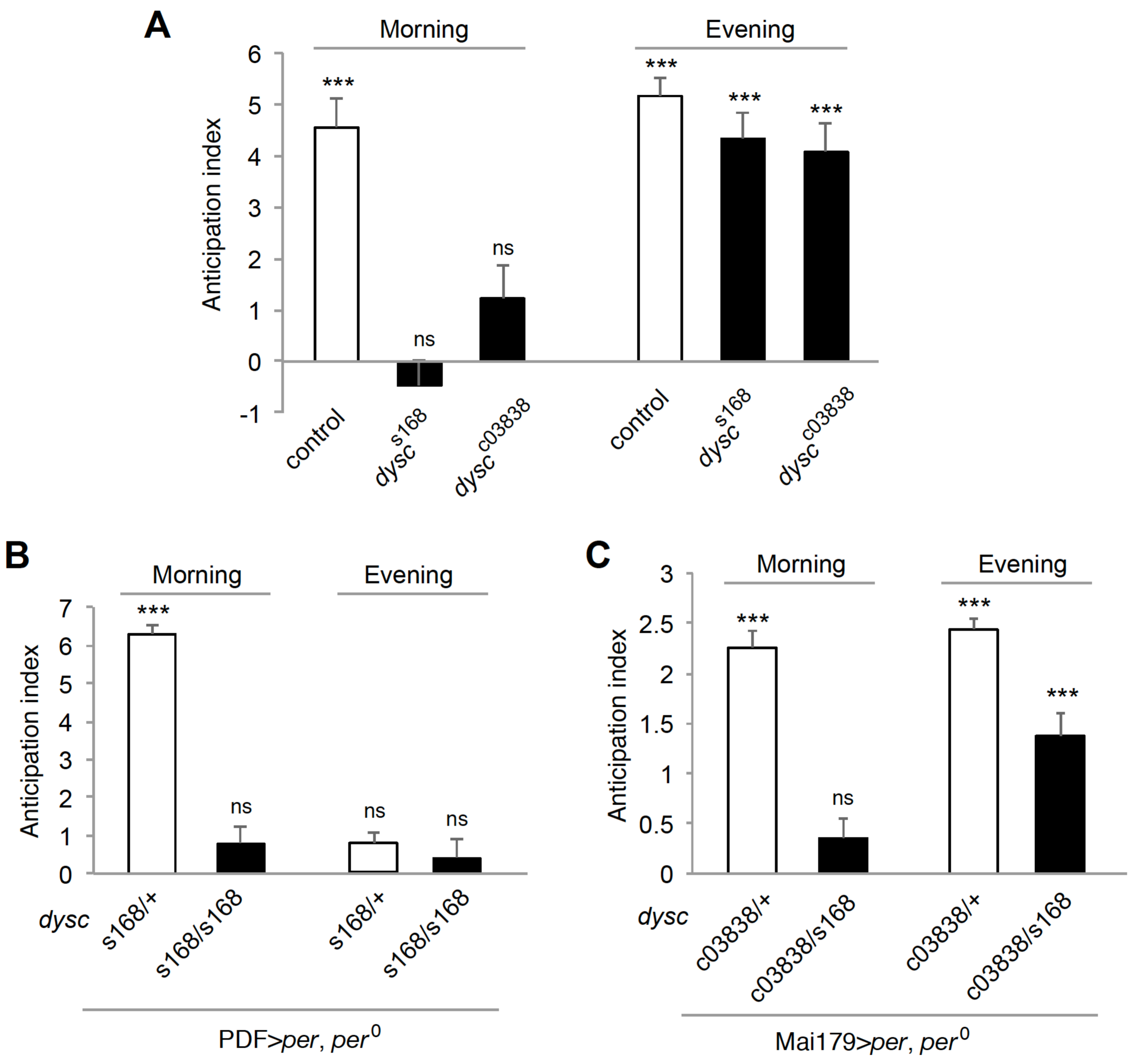


**Figure S1. Loss of DYSC differentially affects morning and evening oscillators.**

**(A-C)** Anticipation index for the data presented in Figures 1A (A), 1C (B), and 1D (C). FaasX was used to calculate anticipation index, defined as the slope of the best-fitting regression line for the activity counts over a period of 4 h (controls) or 8 h (all other genotypes) prior to a light-dark transition. Bars represent anticipation index ± standard error, and statistics indicate whether anticipation index is significantly different from 0. *** $p < 0.0005$, ns: not significant, *t* test with Bonferroni correction.

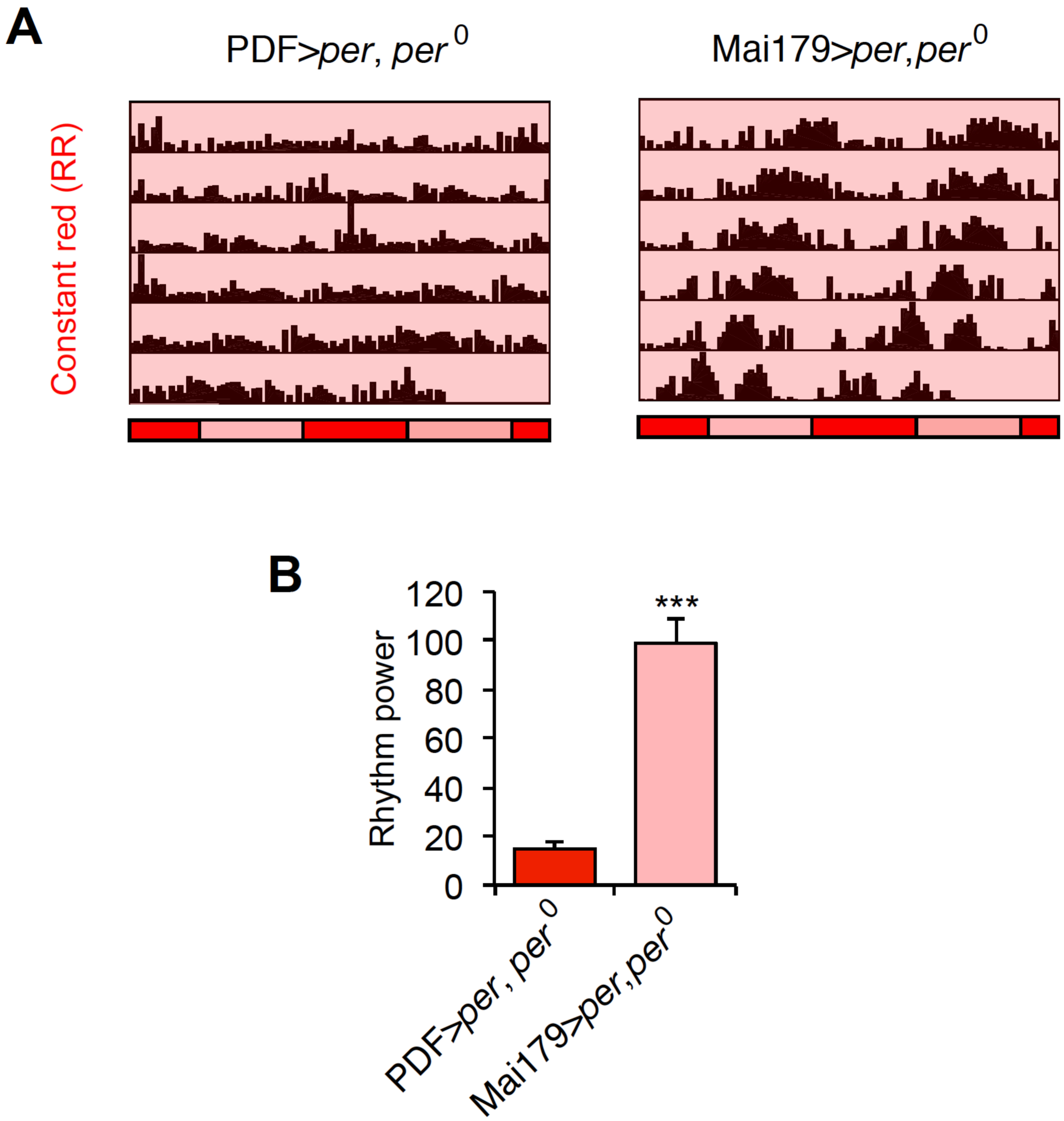


**Figure S2. Rhythmic behavior in RR is driven by the evening oscillator. (A)** Representative actograms of *per*$^0$ flies in which *per* expression is restored in the morning oscillator using PDF-Gal4 or in both morning and evening oscillators using Mai179-Gal4 in RR. **(B)** Mean power of rhythmicity of flies of the same genotypes as in A. n = 31-32. Bars represent mean ± s.e.m. *** $p < 0.0005$, *t* test.

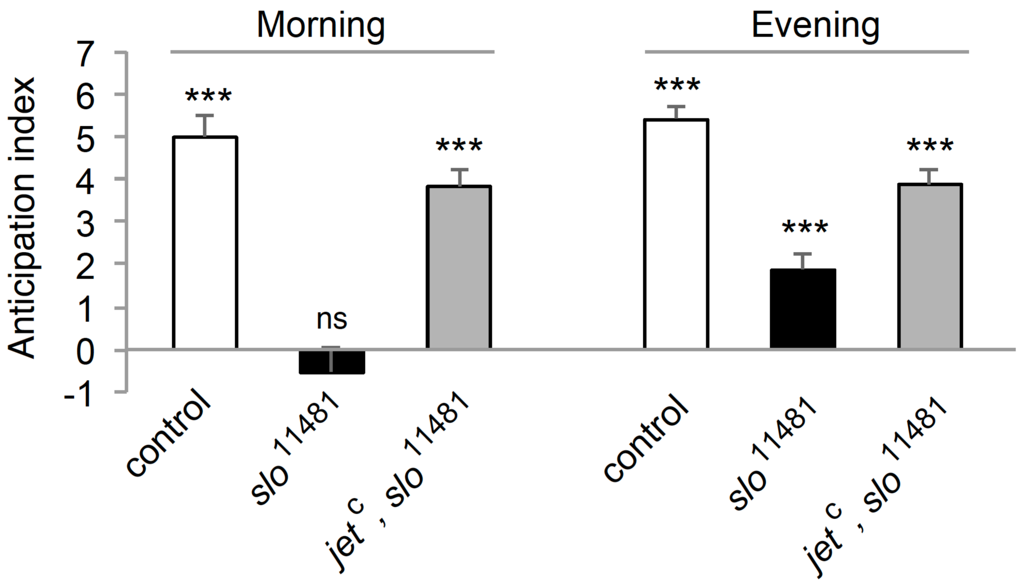


**Figure S3. Reduced JET rescues morning anticipation in *slo* mutants. (A)** Anticipation index for the data presented in Figure 4C. Anticipation index is calculated over a period of 4 h (controls) or 8 h (*slo* mutants) prior to a light-dark transition. Bars represent anticipation index ± standard error, and statistics indicate whether anticipation index is significantly different from 0. *** $p < 0.0005$, ns: not significant, *t* test with Bonferroni correction.
